# Epithelial Regeneration Ability of Crohn’s Disease Assessed Using Patient-Derived Intestinal Organoids

**DOI:** 10.1101/2021.03.25.437111

**Authors:** Chnasu Lee, Sung Noh Hong, Eun Ran Kim, Dong Kyung Chang, Young-Ho Kim

## Abstract

**Background:** The intrinsic limitation of cell lines and animal models limits our understanding of epithelial regeneration capability in Crohn’s disease (CD). Therefore, we aimed to study epithelial regeneration ability in CD using an intestinal organoid model. Further, since tumor necrosis factor alpha (TNFα) is a major proinflammatory effector during CD pathogenesis, we also investigated TNFα-induced alteration of regeneration ability in CD patient-derived intestinal organoids.

**Methods:** Human intestinal organoids were constructed in a three-dimensional intestinal crypt culture of enteroscopic biopsy samples from control subjects and patients with CD. The epithelial regeneration ability of intestinal organoids was assessed using organoid reconstitution, 3-(4,5-dimethylthiazolyl-2)-2,5-diphenyltetrazolium bromide (MTT), 5-ethynyl-2′-deoxyuridine (EdU), and wound healing assays.

**Results:** *Ex vivo* cultures of ileal crypt cells revealed that organoid formation rate of CD patients were reduced compared with that of control subjects (*p* <.001). CD patient-derived organoids sub-cultured for more than 6 passages showed stable organoid reconstitution and identical morphological features. The organoid constitution and MTT assay revealed that the viability of TNFα-treated CD patient-derived organoids were significantly lower than that of TNFα-treated control organoids (*p* <.05 for each). The number of EdU+ proliferative cells was significantly lower in TNFα-treated CD patient-derived organoids than in TNFα-treated control organoids (*p* <.05). The wound-healing ability of TNFα-treated CD patient-derived organoids was significantly lower than that of TNFα-treated control organoids (*p* <.001).

**Conclusions:** The clinical trials are disabled to settle this issue, our results indicated that the epithelial regenerative ability is impaired in patients with CD, especially in TNFα-enriched condition.

## Introduction

The intestinal epithelium, which acts as a frontline defense of the human body, is repetitively injured and continuously regenerated.^1^ Epithelial regeneration depends on the renewal and proliferation of LGR5+ intestinal stem cells (ISCs), which occur in intestinal crypts.^2^ Crohn’s disease (CD) is a chronic relapsing-remitting inflammatory bowel disease (IBD), characterized by cycles of mucosal inflammation and ulceration, followed by regeneration and restoration.^3^ Impaired epithelial regeneration can lead to sustained intestinal inflammation, and could be accompanied by ulcers and complications, such as fibrosis and fistulas.

Nevertheless, the epithelial regeneration capacity of patients with CD has not been evaluated. Clinical trials could not be implemented to evaluate this issue due to ethical reasons and clinical heterogeneity. Classical cell lines and animal model systems have an intrinsic limitation.^4^ Immortalized cell lines consist of a homogenous cellular component and evade cellular senescence due to the presence of certain mutations.^5^ Animal models are difficult to replicate human-specific biological processes.^6^ With the advent of intestinal epithelium-derived organoids, it is possible to cultivate all epithelial cellular components and re-create the functional crypt-villus architecture^7–9^; patient-derived intestinal organoids might be appropriate for studying the regenerative ability of epithelial cells, and might prove to be a useful model for studying human intestinal diseases. This study aimed to evaluate the epithelial regenerative ability of CD patient-derived intestinal organoids, compared to that of control subject-derived intestinal organoids.

Inflammation challenges the epithelial integrity and barrier function. The intestinal epithelium needs to adapt to a multitude of signals in order to perform the complex process of maintenance and restitution of its barrier function.^10^ Tumor necrosis factor alpha (TNFα) acts as a major proinflammatory and tissue damage-promoting effector during the pathogenesis of CD; this is supported by evidence provided by studies involving experimental mouse models and the therapeutic effects of TNFα-neutralizing reagents in IBD treatment.^11–13^ Mucosal healing, which can be regarded as the restitution of the intestinal epithelial lining, is generally defined as a regression of endoscopic lesions in CD. Recently, mucosal healing has been considered as a therapeutic target, as increased mucosal healing improves the prognosis of patients with CD further.^14^ We should attempt to understand the TNFα-induced alteration of epithelial regenerative ability in CD patient-derived organoids, compared to that of control organoids.

## Materials and Methods

### Sampling

To establish intestinal organoids, we used human intestinal tissues obtained from control subjects and CD patients using biopsy forceps during single-balloon enteroscopy at the Samsung Medical Center, Seoul Korea, between November 2016 and December 2018. At least four biopsy samples were obtained from mucosa tissues in the jejunum (100–150 cm distal of the ligamentum of Treitz), ileum (50–100 cm proximal to the ileocecal valve), and colon (transverse colon). Patients with CD were diagnosed according to the practice guidelines.^15^ In patients with CD, biopsies were performed at least 5 cm away from ulcers. All samples were obtained after receiving informed consent. This study was approved by the institutional ethical committee of the Samsung Medical Center (IRB No. 2016-02-022).

### Crypt isolation from biopsies

Endoscopic biopsy samples were incubated in PBS with 10 mM ethylenediaminetetraacetic acid (Thermo Fischer Scientific, San Jose, CA, USA) and 1 mM dithiothreitol (Thermo Fischer Scientific) at 4°C for 30 minutes, and then vortexed for 30–120 seconds. The supernatant containing crypts was filtered through 70-µm cell strainers (Corning, Bedford, MA, USA) and suspended in the basal medium advanced Dulbecco’s modified Eagle’s medium (DMEM)/F12 (Thermo Fischer Scientific) supplemented with antibiotic–antimycotic solution (Thermo Fischer Scientific), HEPES (Thermo Fischer Scientific), GlutaMAX (Thermo Fischer Scientific), N2 (Thermo Fischer Scientific), B27 (Thermo Fischer Scientific), and *N*-acetylcysteine (Sigma-Aldrich, St. Louis, MO, USA)).

### Three-dimensional (3D) intestinal crypt culture

Isolated crypts were resuspended in Matrigel (Corning) and plated in 48-well culture plates (Corning). After incubation at 37°C for 15 min, 250 µl of maintenance medium (50% Wnt3a-conditioned medium (ATCC#CRL-2647, Manassas, VA, USA) and 50% of 2× basal medium supplemented with recombinant human EGF (Sigma-Aldrich), recombinant human noggin (R&D Systems, Minneapolis, MN, USA), recombinant human R-spondin1 (PeproTech, Cranbury, NJ, USA), nicotinamide (Sigma-Aldrich), p160ROCK inhibitor (Selleck Chemicals, Houston, TX, USA), p38 MAP kinase inhibitor (SB202190, Sigma-Aldrich), and Prostaglandin E2 (PGE2, Cayman Chemical, Ann Arbor, MI, USA) was added to the wells. A GSK3 inhibitor (Stemgent, Cambridge, MA, USA) was added to the medium for the first 2 days.

Depending on the shape, organoids can be classified into spheroids and enteroids. Spheroids are defined as round- or oval-shaped organoids with a thin wall consisting of a single layer of undifferentiated cells. Enteroids are defined by the presence of visually sharp borders (buddings) along their basolateral (anti-luminal) side or irregular, thickened walls, which consist of all components of epithelial cells, including differentiated and undifferentiated cells.^16^

### Organoid subculture and maintenance

After 7 days of the culture process, organoids in Matrigel were mechanically disrupted and resuspended in cell dissociation buffer (Thermo Fischer Scientific). Single cells and small cell clusters were resuspended in Matrigel and plated in 48-well culture plates (Corning). The medium was changed every 2 days and organoids were passaged using a ratio of 1:2–1:4 on day 7. After 6 passages, most organoids in the maintenance media were uniform spheroids and able to passage stably for a long time.

### Differentiation of intestinal organoids

To recreate the physiological parameters of organs, spheroids were cultured in a differentiation medium (maintenance medium without Wnt3A conditional medium, SB202190, nicotinamide, and PGE2). The differentiation medium was changed every 2 days and enteroids were cultured for 7–12 days.

### Organoid formation assay

One hundred crypts obtained from control subjects and CD patients were plated in 25 µL of Matrigel in maintenance medium. The organoid formation rate was calculated as the percentage of viable organoids per 100 intestinal crypts.

### Organoid reconstitution assay

After more than six passages, intestinal organoids were cultured in maintenance medium for 2 days to obtain a stable number of organoids before inducing organoid differentiation; then, the maintenance medium was replaced with differentiation medium every 2 days. Different concentrations of human recombinant TNFα (R&D Systems) were added to the culture medium every 24 hours. The organoid reconstitution rate was calculated as a percentage of the number of viable organoids on day 9–10 proportional to the number of viable cells on day 2.

### 3-(4,5-Dimethylthiazolyl-2)-2, 5-Diphenyltetrazolium Bromide (MTT) assay

Ten microliters of MTT (Sigma-Aldrich) was added to each well of the culture plates incubated for 3 h until purple precipitate was visible. After the addition of 100 µl of detergent reagent, the organoids were incubated at room temperature in the dark for 2 h, prior to recording the absorbance at 570 nm.

### 5-Ethynyl-2′-Deoxyuridine (EdU) assay

Two hundred micrograms of EdU (Abcam, Cambridge, UK) was added to the culture medium 2 h before fixation with cold 4% paraformaldehyde (Biosesang. Seongnam-si, South Korea). EdU incorporation into DNA was detected using the Click-iT^™^ EdU Alexa Fluor^®^ 488 imaging kit (Thermo Fischer Scientific).

### Wound Healing Assay

Organoids subjected to 3D culture were digested into single cells using TrypLE Express (Thermo Fischer Scientific), and 5 ×10^4^ cells were seeded into 24-well plates containing CytoSelect^™^ 24-Well Wound Healing Assay inserts (Cell Biolabs, San Diego, CA, USA) (*16*). Organoid monolayers were cultured in maintenance medium until confluency was achieved. The inserts were then carefully removed to generate 0.9-mm-diameter wounds and fresh differentiation medium was added to each well. The non-healed wound area was measured in three different areas.

### RNA sequencing

RNA sequencing was performed using total RNA samples containing >10 µg RNA and an integrity number >8. Libraries were constructed for performing whole-transcriptome sequencing using the TruSeq RNA Sample Preparation Kit v2 (Illumina, San Diego, CA, USA) and sequenced using the 100-bp paired-end mode of the TruSeq Rapid PE Cluster Kit and the TruSeq Rapid SBS Kit (Illumina).

### Real-Time Quantitative Reverse Transcription Polymerase Chain Reaction (qPCR)

Total RNA was extracted from intestinal organoids using the RNeasy Mini Kit (QIAGEN, Hilden, Germany). One-step qPCR was performed using One Step PrimeScript^™^ III RT-qPCR Mix (Takara, Kusatsu, Japan) with primers as follows; LGR5 primer (Forward: 5’-aactttggcattgtggaagg-3’, Reverse: 5’-acacattgggggtaggaaca-3’), BMI1 primer (Forward: 5’-cgtgtattgttcgttacctgga-3’, Reverse: 5’-ttcagtagtggtctggtcttgt-3’), ATOH1 primer (Forward: 5’-cagctgcgcaatgttatccc-3’, Reverse: 5’-ttgtagcagctcggacaagg-3’), and HES1 primer (Forward: 5’-tttcctcattcccaacgggg-3’, Reverse: 5’-ctggaaggtgacactgcgtt-3’).

## Results

To evaluate the organoid formation ability of intestinal crypts, intestinal crypts isolated from duodenal (CD, n = 5; controls, n = 5), jejunal (CD, n = 9; controls, n = 13), ileal (CD, n = 43; controls, n = 15), and colonic samples from (CD, n = 12; controls, n = 8) control subjects (n = 34) and CD patients (n = 51) were cultured in Matrigel in maintenance medium (Figure2). Characteristics of the enrolled patients are listed Table 1. In both groups, the organoid formation rate of duodenal crypts was the highest, followed by jejunal, ileal, and colonic crypts. The organoid formation rate of the ileal crypts obtained from patients with CD was significantly lower than that of ileal crypts obtained from controls at day 3 (53.4% ± 4.0% *vs*. 39.6% ± 2.4%, *p* = 0.005), day 5 (38.9% ± 3.0% *vs*. 21.1% ± 1.7%, *p* < 0.001), and day 7 (29.2% ± 2.8% *vs*. 15.7% ± 1.5%, *p* < 0.001). The organoid formation rate of jejunal crypts obtained from patients with CD was numerically lower than that of those obtained from controls (46.8 ± 2.6% *vs*. 38.1% ± 3.8%) (Figure 2).

**Figure 1.**
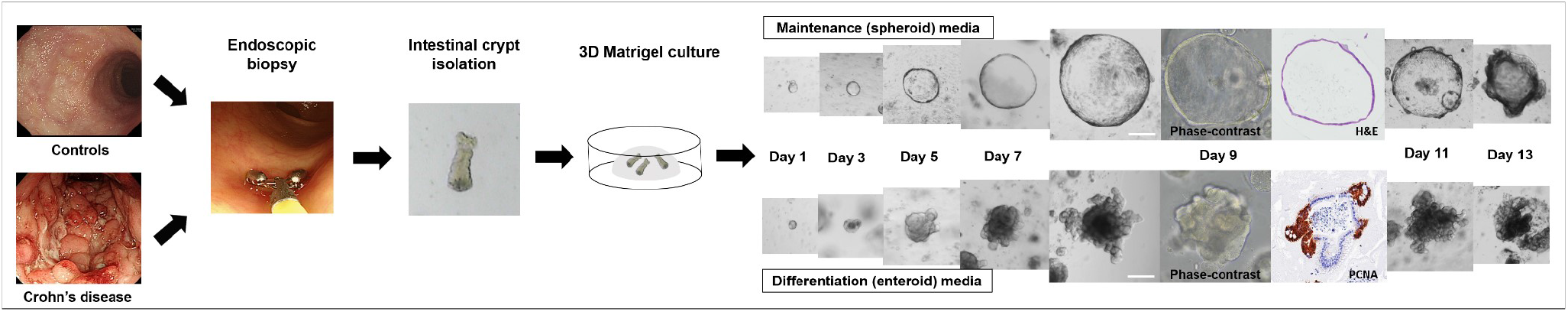
Flow diagram of the patient-derived intestinal organoid model. White scale bar = 200 µm.

**Figure 2.**
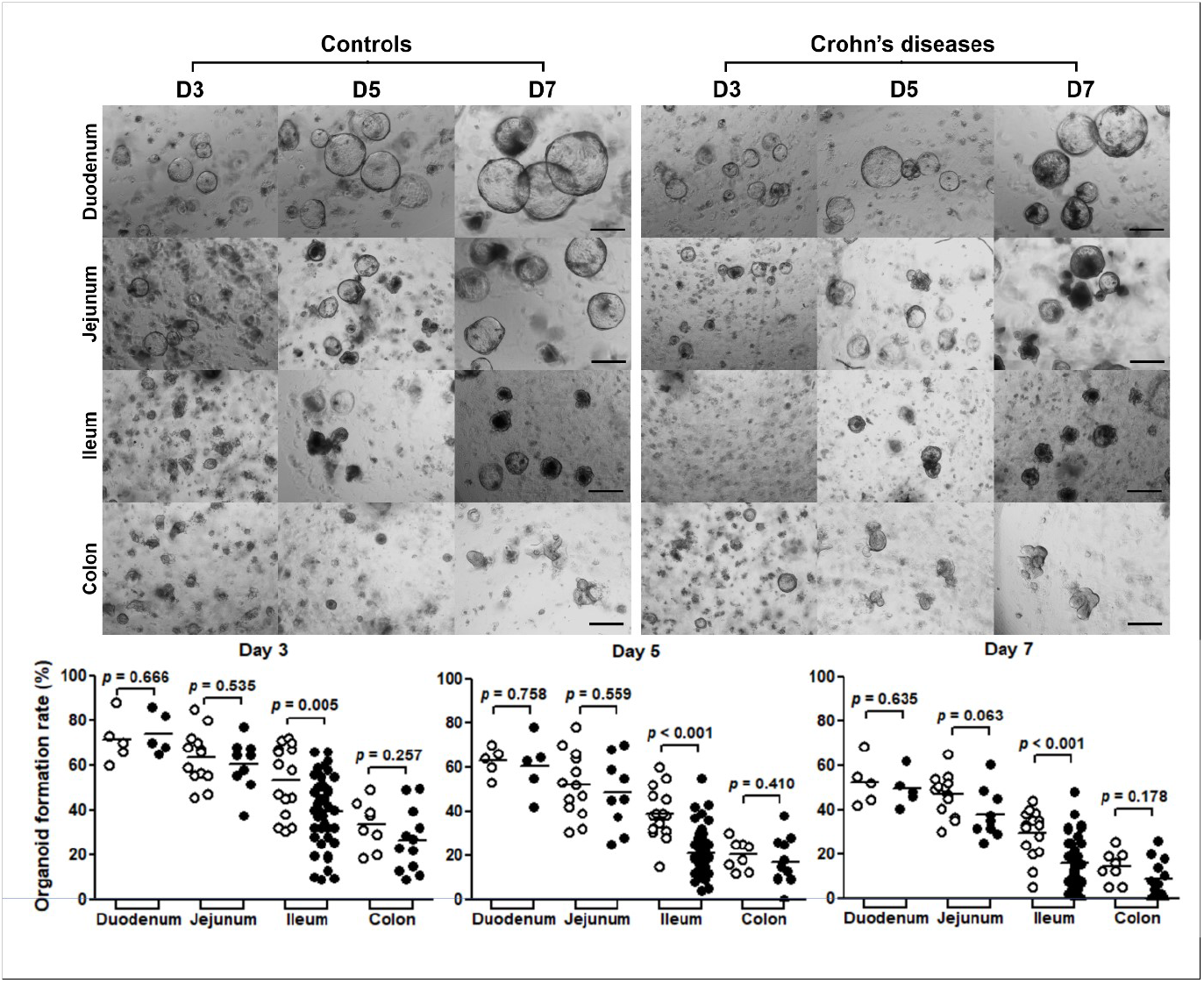
Organoid formation assay. Intestinal crypts were isolated from the duodenum (n=5), jejunum (n=9), ileum (n=43), and colon (n=12) of patients with CD and the duodenum (n=5), jejunum (n=13), ileum (n=15), and colon (n=8) of controls. Differences of organoid formation between the controls and patients with Crohn’s diseases in each segment of intestine were evaluated by *t*-test; black scale bar = 500 µm.

**Table 1.** Characteristics of enrolled patients and number of attempts to construct organoids.

|  |  |  |  |
| --- | --- | --- | --- |
| Total number of enrolled patients | 85 | Number of attempts to construct organoids | 110* |
| Patients with CD (n) | 51 | Number of sub-culture (≥ passage 3) enabled organoids | 61 |
| Age, years (mean ± S.D) | 36.2 ± 13.4 | CD-derived organoids (n) | 69 |
| Male (n) | 43 | Duodenal organoids (n of attempts / n of subculture) | 5 / 4 |
| Montreal classification (n) |  | Mucosal healing / no duodenal involvement | 5 / 4 |
| Age at diagnosis (A1/A2/A3) | 0 / 42 / 9 | Active ulcer | 0 / 0 |
| Location (L1/L2/L3) | 1 / 34 / 16 | Jejunum organoids (n of attempts / n of subculture) | 9 / 8 |
| Behavior (B1/B2/B3) | 20 / 27 / 4 | Mucosal healing / no jejunal involvement | 4 / 4 |
| Medication at sampling |  | Active ulcer | 5 / 4 |
| None (sampling before treatment), (n) | 6 | Ileal organoids (n of attempts / n of subculture) | 43 / 23 |
| Immunomodulator user, (n) | 25 | Mucosal healing / no ileal involvement | 8 / 5 |
| Anti-TNFα user, (n) | 25 | Active ulcer | 35 / 18 |
| Sampling modality |  | Colonic organoids (n of attempts / n of subculture) | 12 / 2 |
| Colonoscopy | 11 | Mucosal healing / no colonic involvement | 4 / 1 |
| Single-balloon enteroscopy (n) | 40 | Active ulcer | 8 / 1 |
| Controls (n) | 34 | Control-derived organoids† | 41 / 24 |
| Age, years (mean ± S.D) | 50.6 ± 17.9 | Duodenal organoids (n of attempts / n of subculture) | 5 / 4 |
| Male sex (n) | 24 | Jejunum organoids (n of attempts / n of subculture) | 13 / 11 |
|  |  | Ileal organoids (n of attempts / n of subculture) | 15 / 8 |
|  |  | Colonic organoids (n of attempts / n of subculture) | 8 / 1 |

\*patients having samples from two (n = 21) and three (n = 2) different sites.

Organoids grown from intestinal crypts were sub-cultured in maintenance medium and the control organoids became uniformly spheroid (> 90% of total organoids) after 2–4 passages. However, CD patient-derived organoids exhibited both enteroid and spheroid shapes in the early passages and became uniformly spheroid after 4–6 passages. After six passages, control and CD patient-derived organoids showed similar stable morphological features (Figure 3A). The intestinal epithelial cell lineage-specific gene expression levels of control and CD patient-derived organoids were similar at the sixth passage. The expression of differentiated epithelial cell markers, such as enterocytes (ECs), goblet cells (GCs), Paneth cells (PCs), and enteroendocrine cells (EECs) increased with time, whereas the ISC marker was stably expressed (Figure 3B).

**Figure 3.**
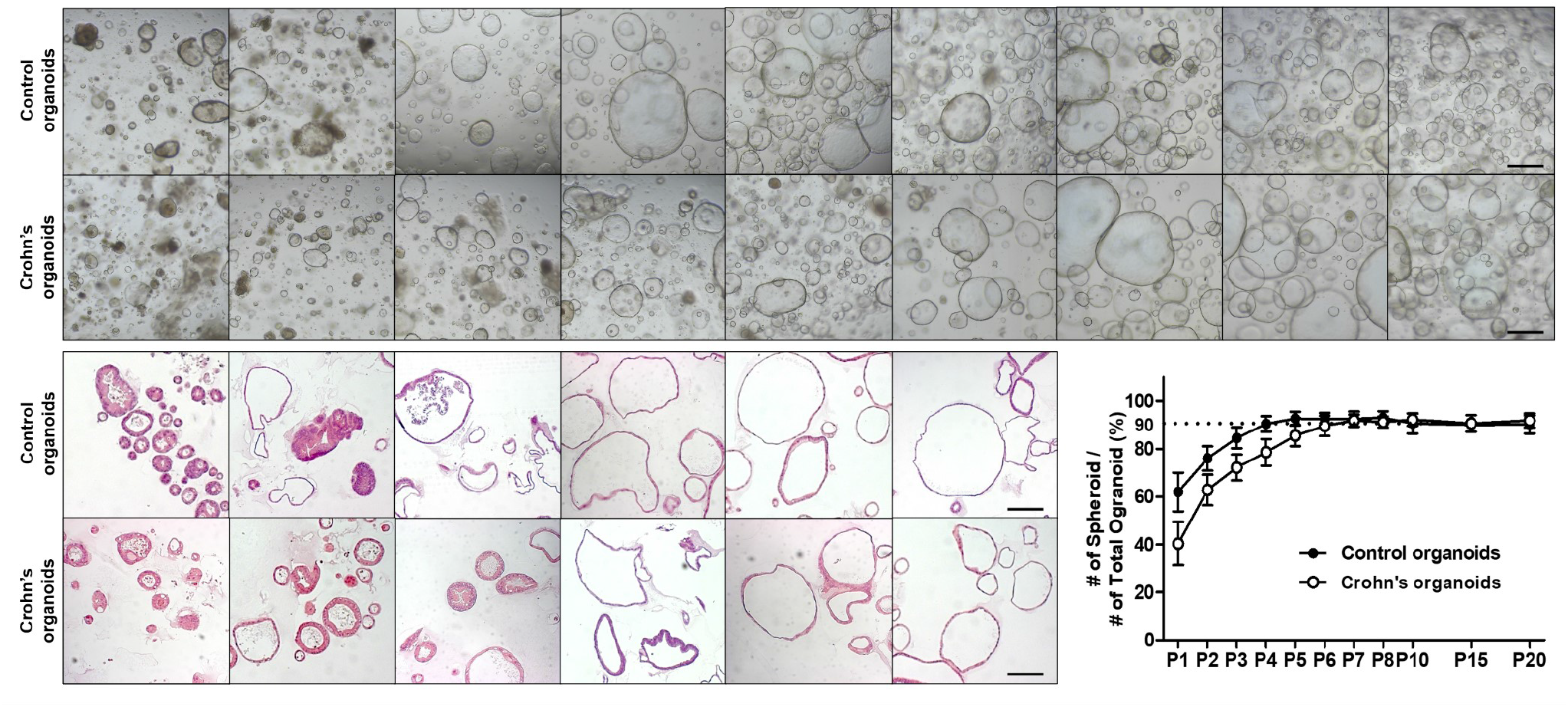
Long-term culture of control and CD patient-derived organoids. (A) Bright field and H&E staining images of long-term culture of control and CD patient-derived organoids. After six passages, the morphology of control and CD patient-derived organoids became identical. Black scale bar = 500 µm. (B) Expression pattern of epithelial cell markers of control and CD patient-derived organoids at the sixth passage. The expression of intestinal epithelial cell lineage-specific genes after the sixth passage in the control and CD patient-derived organoids cultured for 3, 6, and 9 days showed a similar pattern (n=2 for each).

The epithelial regenerative ability of intestinal organoids was assessed via organoid reconstitution, wound healing assays, and the use of 3-(4,5-dimethylthiazolyl-2)-2,5-diphenyltetrazolium bromide (MTT) and 5-ethynyl-2′-deoxyuridine (EdU). The shapes of organoids cultured in the maintenance medium were similar regardless of location; however, those cultured in differentiation medium tended to have different shapes, depending on their origin. Ileal organoids typically exhibited budding, whereas jejunal organoids formed thick-walled structures (Supplementary Figure S1). The cell viability—measured using MTT and the enteroid/spheroid ratio—decreased significantly with an incremental increase in the TNFα concentration. At TNFα concentrations of ≤ 10 ng/ml in the differentiation medium, changes in cell viabilities and morphologies were negligible; however, the changes observed at TNFα concentrations of ≥ 30 ng/ml were remarkable. In addition, organoids became senescent by day 9–10 (Supplementary Figures S2) and changes in *LGR5, BMI1, HES1*, and *ATOH1* expression levels occurred within 24 h after TNFα treatment (Supplementary Figure S3).

Based on these results, control and CD patient-derived organoids —cultured over the long-term (passage ≥ 6)—were treated with 30 ng/ml TNFα every 24 hours for 10 days (Figure 4A). The organoid reconstitution rate of TNFα-treated organoids was significantly lower than that of TNFα-free organoids (*p* < 0.001). There was no significant difference in the organoid reconstitution rate between TNFα-free controls and CD patient-derived organoids; however, the organoid reconstitution rate of TNFα-treated CD patient-derived organoids was significantly lower than that of TNFα-treated control organoids (*p* < 0.05 for both jejunal and ileal organoids; Figures 4B and 4C).

**Figure 4.**
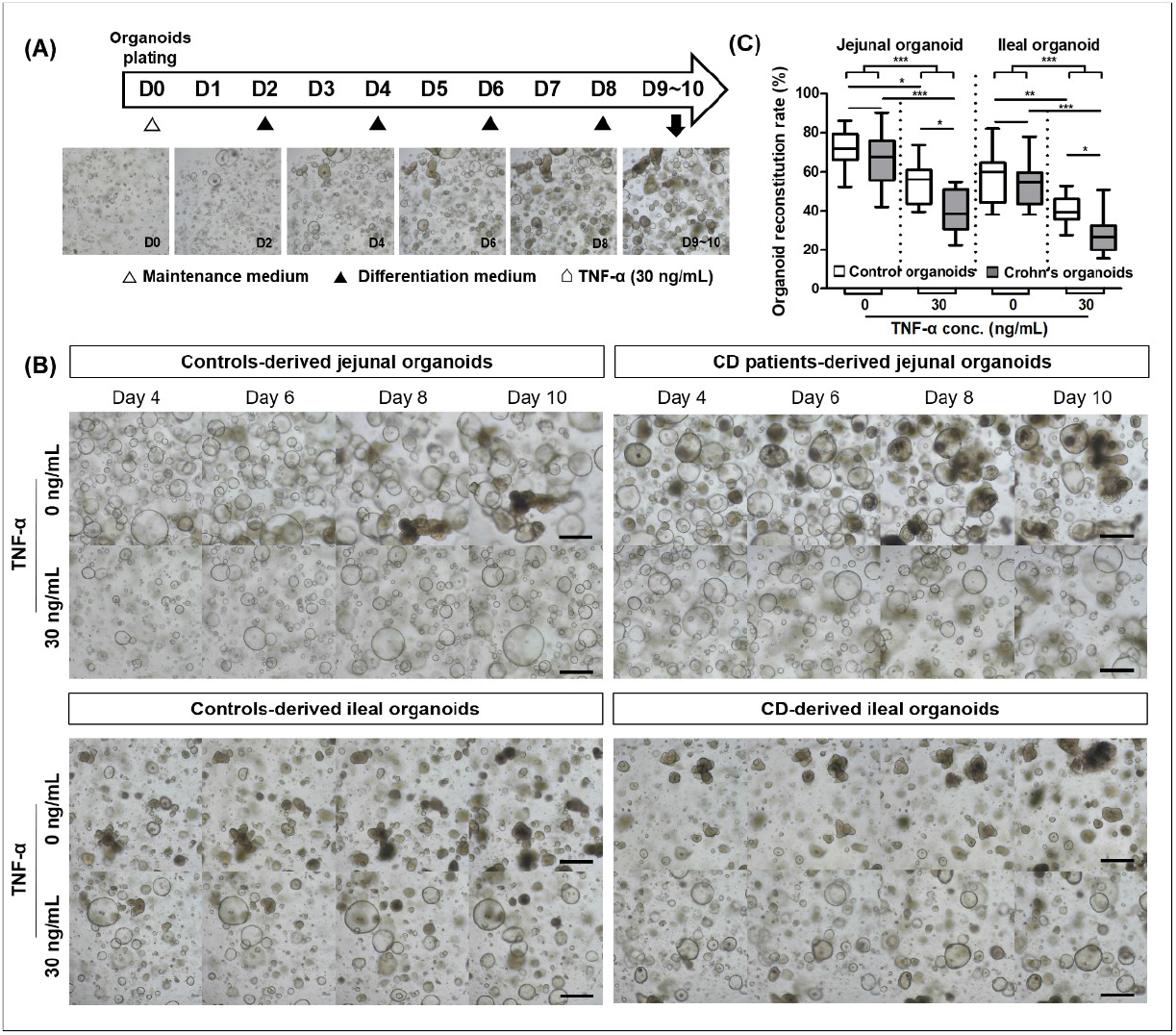
Organoid reconstitution assay in tumor necrosis factor-alpha (TNFα) enriched conditions. (A) Study flow diagram. (B) Organoid reconstitution assay of jejunal and ileal control and CD patient-derived organoids (n=5 each) in TNFα-enriched conditions. Assays were conducted in triplicate. (C) Organoid reconstitution rate. Reconstituted organoid number is expressed as a percentage value, based on values of TNFα-free control jejunal organoids. Differences were evaluated using ANOVA with Bonferroni’s multiple comparison test; \**p* <0.05, \*\**p* <0.01, and \*\*\**p* <0.001.

The organoid viability was assessed using MTT; results showed that the formazan absorbance values of viable TNFα-treated organoids were significantly lower than those of TNFα-free organoids (*p* < 0.001 for both jejunal and ileal organoids). In the TNFα-enriched condition, the number of viable jejunal and ileal CD patient-derived organoids was significantly lower than that in the control organoids (*p* < 0.05 for both jejunal and ileal organoids; Figure 5A). Two hours after EdU administration, EdU+ cells were confirmed in the buds, which represented the intestinal crypt (Supplementary Figure S4). The number of EdU+ cells was higher in TNFα-free organoids than in TNFα-treated organoids (*p* < 0.001). Although there was no significant difference in the number of EdU+ cells between the control and CD patient-derived organoids in the steady state, the number of EdU+ cells were significantly lower in TNFα-treated CD patient-derived organoids than in TNFα-treated control organoids (*p* < 0.05, Figure 5B).

**Figure 5.**
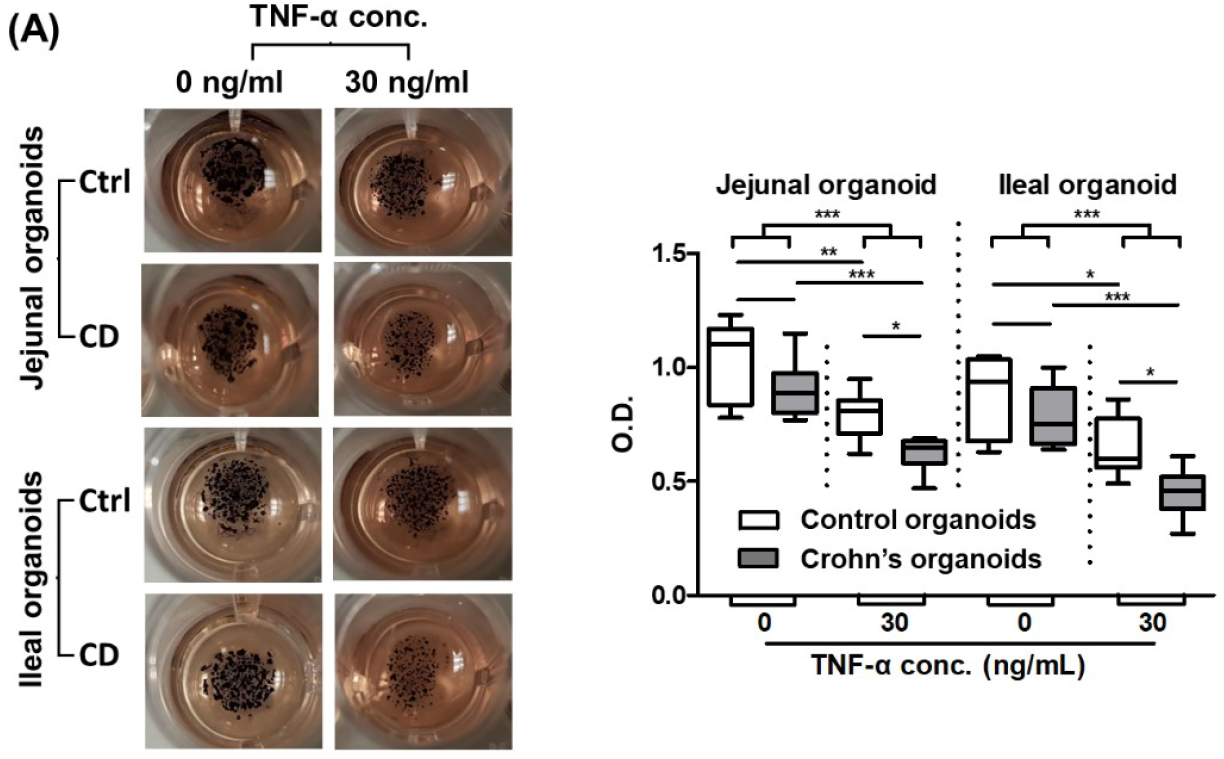
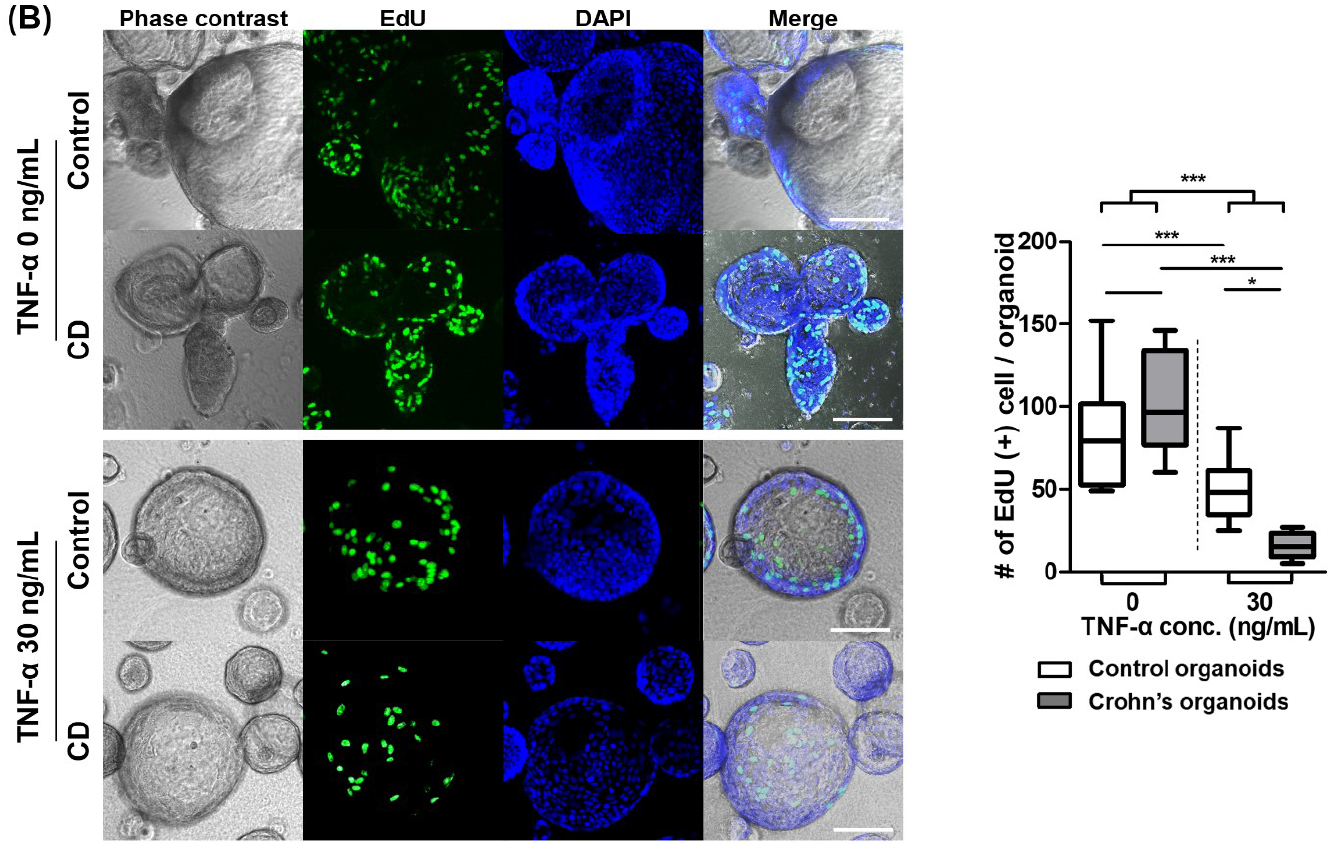
Organoid viability ad proliferation assay in TNFα enriched conditions. (A) 3-(4, 5-dimethylthiazolyl-2)-2, 5-diphenyltetrazolium bromide (MTT) assays. MTT assay were performed in triplicate using jejunal and ileal control and CD patient-derived organoids (n=3 each). (B) 5-ethynyl-2′-deoxyuridine (EdU) assay. EdU+ cell number was measured 2 h after EdU administration into 10 organoids from TNFα-treated and -free control and CD patient-derived organoids (n=3 each). Differences were evaluated using ANOVA and the Bonferroni’s multiple comparison test; \**p* <0.05, \*\**p* <0.01, and \*\*\**p* <0.001.

The wound healing assay showed that the unhealed wound area in TNFα-treated CD patient-derived organoids was significantly larger than that in TNFα-treated control organoids, 16 and 24 h after insert removal (*p* < 0.001 for each). The wound-healing ability of TNFα-treated CD patient-derived organoids was significantly lower than that of TNFα-treated control organoids (*p* < 0.001, Figure 6).

**Figure 6.**
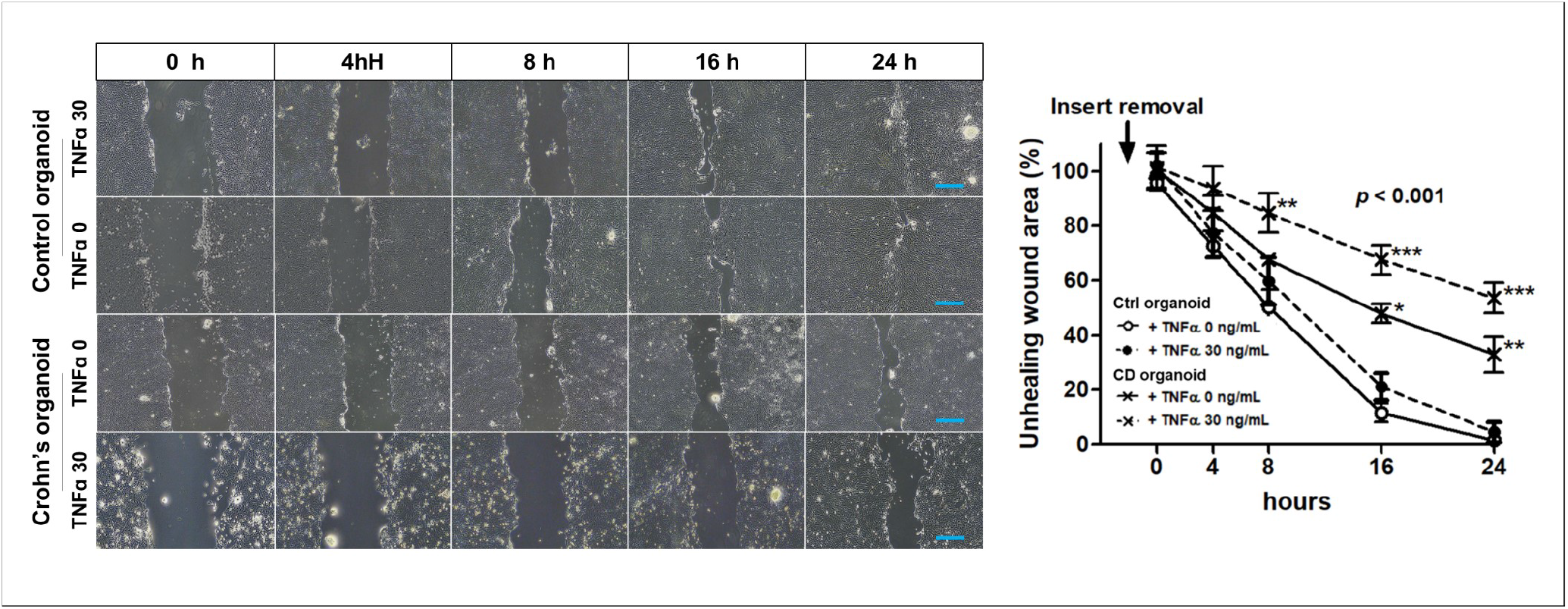
Wound healing assay. Non-healing wound areas in three different areas selected from TNFα treated control and CD patient-derived organoids were measured (n=3 each). Differences were evaluated using the two-way ANOVA test with Bonferroni’s multiple comparison test. Blue scale bar = 4 mm.

## Discussion

Intestinal epithelium physically separates the microbially dense intestinal lumen from the sterile sub-mucosa, which includes structures such as the lamina propria, lymphatic capillary-lacteals, and resident immune cells.^17^ The disruption of the intestinal epithelial lining can provide altered immunologic responses against the recognition of commensal gut microbiota, leading to inflammation and loss of homeostasis in the mucosa.^18^ The rapid restoration of epithelial defects following injuries or physiologic damage is indispensable for maintaining gut homeostasis.^10^ CD is characterized by a chronic and transmural inflammation that can be occurred along the entire gastrointestinal tract, from mouth to anus, but primarily occurs within the small intestine. Nevertheless, the ability of epithelial regeneration and wound healing in patients with CD have not been evaluated, especially under proinflammatory cytokine-enriched conditions. To our knowledge, this study is the first to identify the epithelial regenerative ability is reduced in CD using patient-derived organoid models.

Several previous observations have suggested the impaired ability of epithelial regeneration and wound healing in patients with CD. Patients with CD have a higher post-operative complication rate, especially anastomotic complications leading to intra-abdominal sepsis, than patients without an inflammatory state.^19^ Exposure to ionizing irradiation provokes cell death of rapid proliferating cells, leading acute or chronic gastrointestinal (GI) toxicity in a dose- and time-dependent manner.^20^ Patients with CD experienced an increased risk of GI toxicities following exposure to therapeutic doses of ionizing radiation.^21^ The mutation of CD susceptibility genes, such as NOD2 and ATG16L1, were associated with the dysfunction of ISCs, contributing the impaired cellular regeneration.^22, 23^ Single cell RNA-seq identified the alterations of ISC properties in CD patient-derived small intestinal organoids.^24^

This study identified that the organoid formation ability of CD patient-derived ileal crypts was significantly impaired; this was correlated with the fact that the ileum was the most frequently affected area in CD.^25^ These ileal crypts were endoscopically obtained from the uninflamed intestine; however, the effect of pre-existing microscopic inflammation on organoid formation cannot be completely excluded. Long-term cultured CD patient-derived organoids were passaged in conditions deprived of inflammation and showed no significant differences in the morphology, histology, and gene expression, compared to control organoids, which may mimic the “deep remission” mucosal and histological healing state. Deep remission is considered to be an ideal therapeutic target for the long-term remission of IBD.^3^ However, despite patients achieving deep remission, triggers such as such as inflammatory cytokines and non-steroidal anti-inflammatory drugs (NSAIDs) might induce mucosal inflammation recurrence and IBD ulceration. In this study, TNF was used as a trigger to mimic the inflammatory milieu. TNFα reduced the cell viability, proliferation, wound healing, and organoid formation capacity of intestinal organoids, which was more remarkable in CD patient-derived organoids compared with control organoids. In practice, impaired epithelial regeneration in patients with CD is due to the defective mucosal integrity and sustained intestinal inflammation, leading to ulcers, fibrosis, and fistula, which are the main indication for surgery in patients with CD.26

Previous studies have found that wound healing was accomplished by epithelial restitution, ISC proliferation, and differentiation.^27–29^ Although these wound-healing processes overlapped,, we assessed epithelial restitution by performing analyses using wound healing assays and achieving organoid cell proliferation using organoid reconstitution, MTT, and the EdU assay. In addition, following injuries, wound healing is regulated by a broad spectrum of regulatory factors, including cytokines, growth factors, adhesion molecules, and phospholipids ^10, 30^. Inflammatory processes might especially interfere with epithelial cell migration and proliferation, and thus modulate intestinal epithelial healing.

The limitation of our study is that the organoid culture system does not reflect the effects of intestinal microbiota, dietary components, and the mucosal immune system. The intestinal microbiota and dietary components contribute to the fine-tuning of ISC survival and differentiation.^31^ In addition, only ileal crypts of CD showed a significant reduction in the organoid formation rate compared with those of controls. Because enteroscopy was usually performed for the diagnosis and treatment of CD, ileal sampling was relatively convenient and small sample size of duodenal and jejunum was small. The lack of significance of duodenal and jejunal organoid reconstitution rate might be attributed to small sample size.

Epithelial regeneration depends on the renewal and proliferation of LGR5+ ISCs. Impaired epithelial regeneration can lead to sustained intestinal inflammation. Organoid formation ability of ileal crypt in patients with CD was reduced compared with those of control subjects. When TNFα was used as a trigger to mimic the inflammatory milieu, the epithelial regeneration ability of intestinal organoids was more remarkable in CD patient-derived organoids, compared to that in control organoids. The clinical trials are disabled to settle this issue due to ethical reasons and clinical heterogeneity, our results indicated that the epithelial regenerative ability is impaired in patients with CD, especially in TNFα-enriched condition. In practice, impaired epithelial regeneration in patients with CD induces the defective mucosal integrity and sustained intestinal inflammation, leading to ulcers, fibrosis, and fistulas, which are the main indications for surgery in patients with CD. ^26^ Even though anti-TNFα agents are most effective therapeutic option of CD, around 10–30% of patients do not respond to anti-TNFα agents and 30–50% of patients lose response over time, resulting in complications, hospitalization, and surgery.^32^ To improve this unmet needs, additional treatment option to promote mucosal healing should be explored.

## Acknowledgments

I would like to thank Prof. Martín G. Martín, (Department of Pediatrics, Mattel Children’s Hospital and the David Geffen School of Medicine, University of California Los Angeles), Prof. Matthias Stelzner (Department of Surgery, Veterans Affairs Greater Los Angeles Healthcare System), and Prof. James C Y Dunn (Division of Pediatric Surgery, Department of Surgery, Stanford University) who taught and trained me in construct intestinal organoid models, despite of their busy schedules.

## Supplemental Figure Legends

**Supplementary Figure S1.**
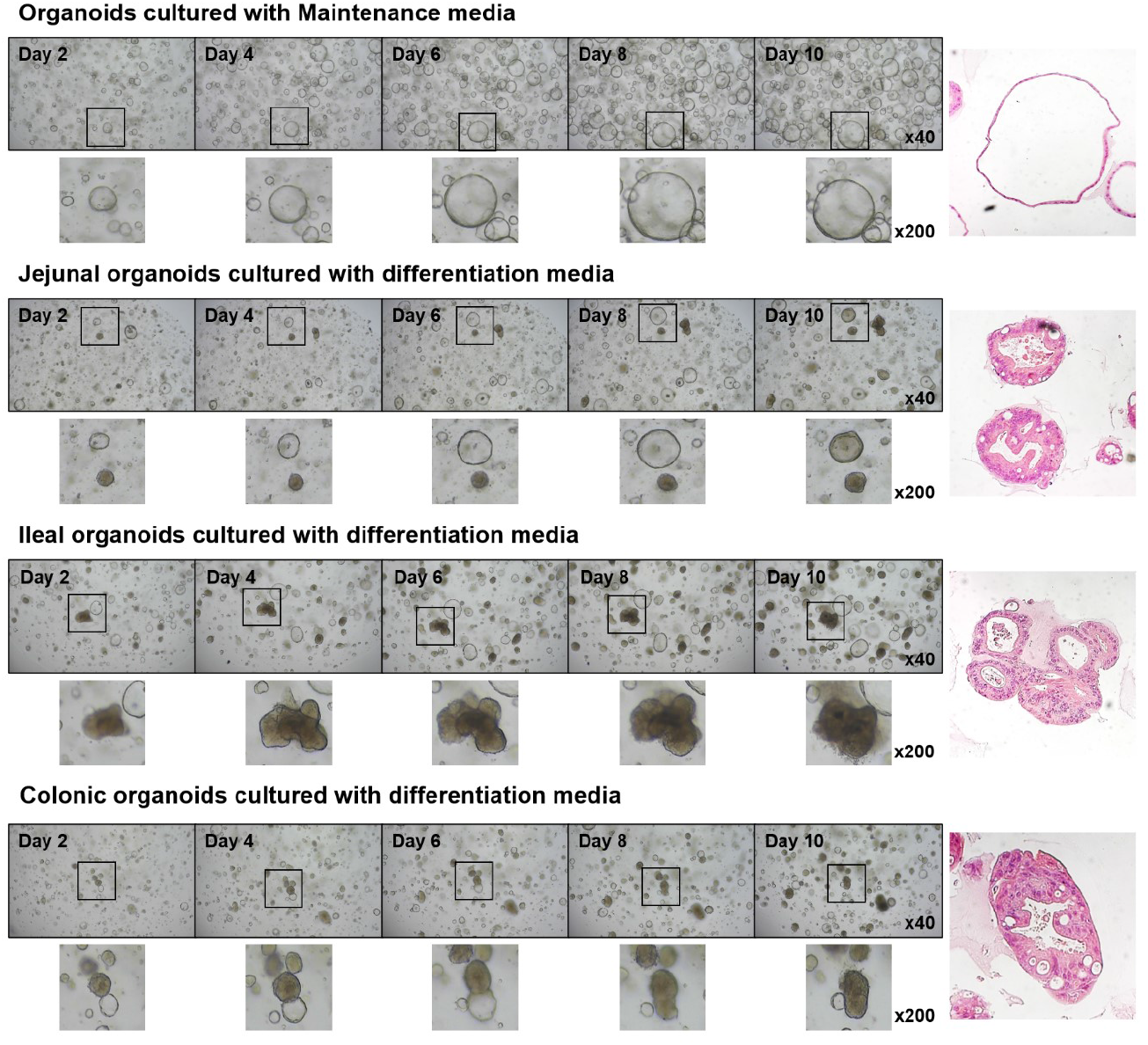
Human intestinal organoid responses to different conditions. Shapes of control intestinal organoids, depending on their intestinal location of origin. Control organoids cultured in maintenance medium were spheroid, whereas those cultured in differentiation medium were enteroid in shape.

**Supplementary Figure S2.**
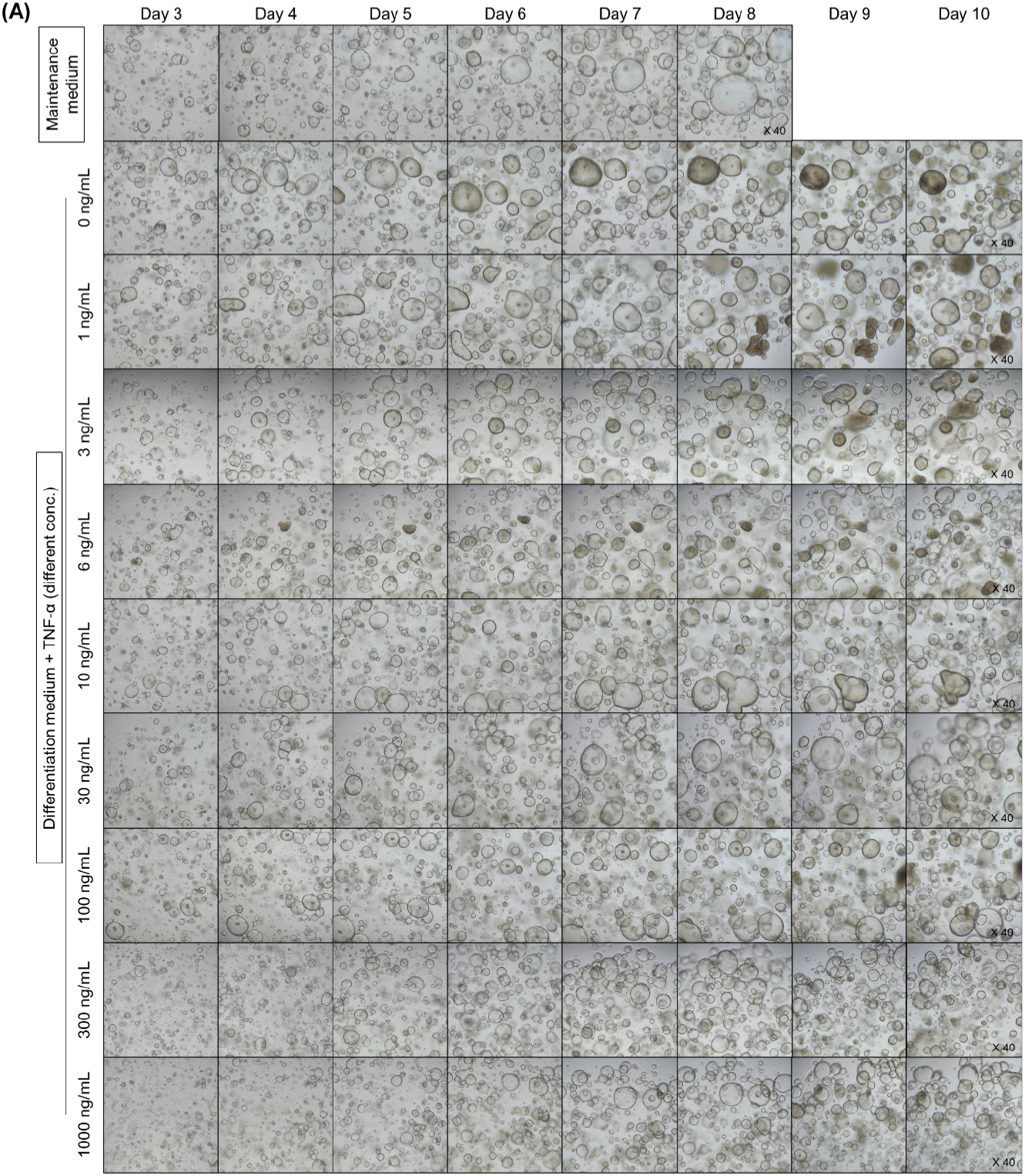
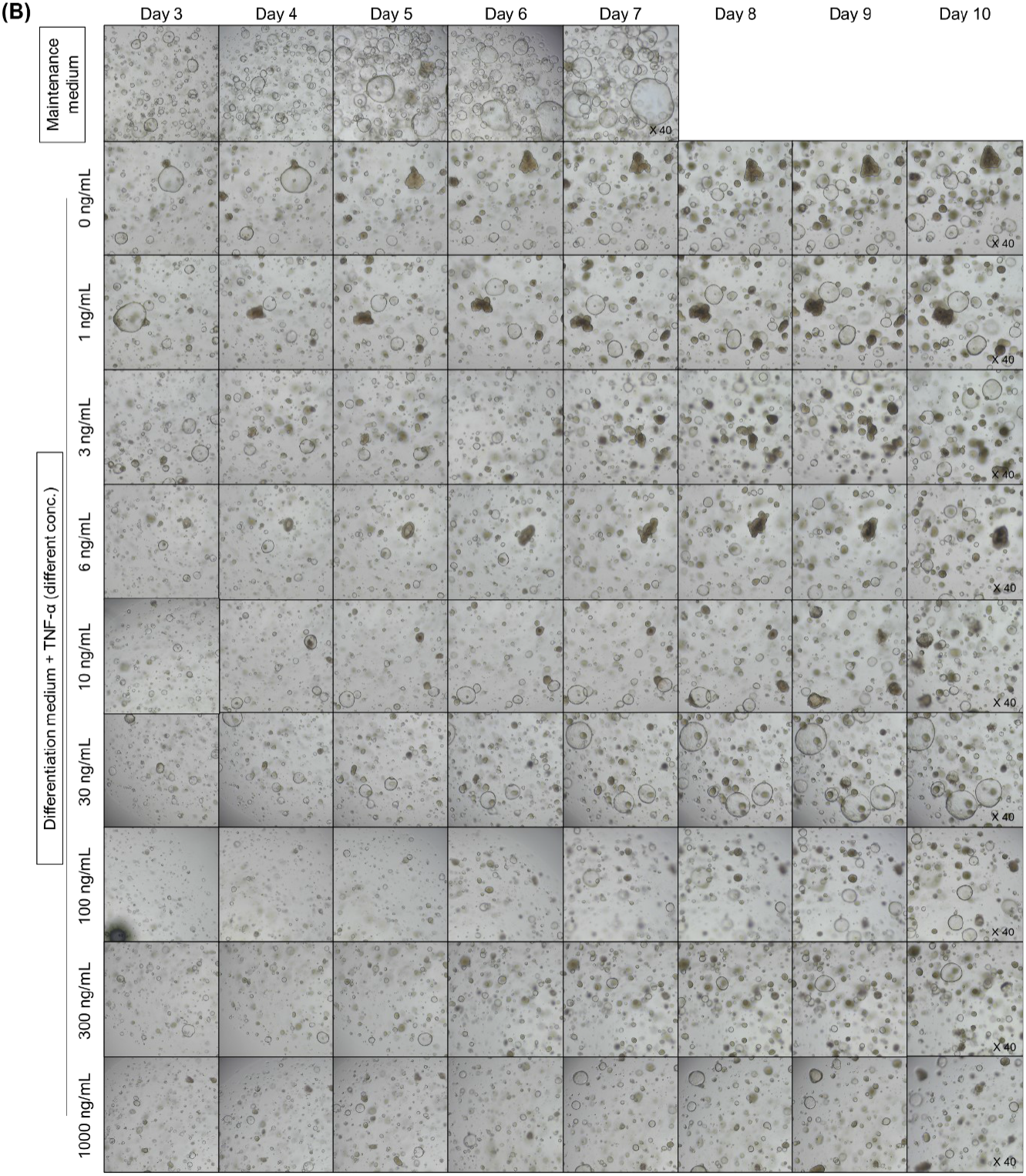
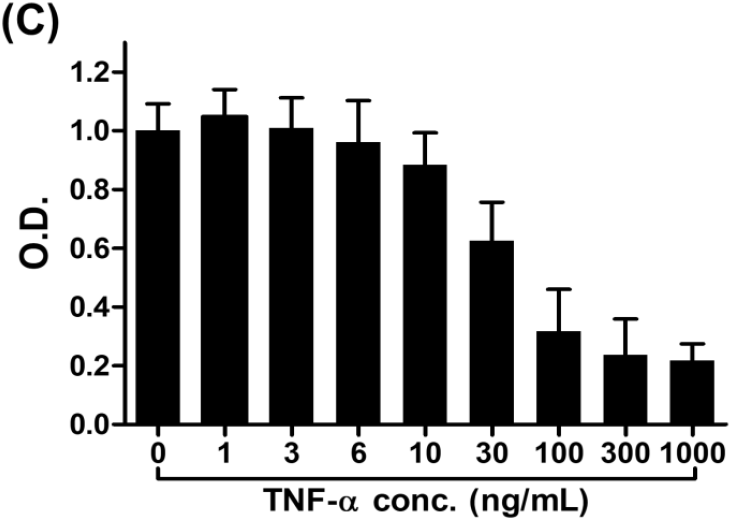
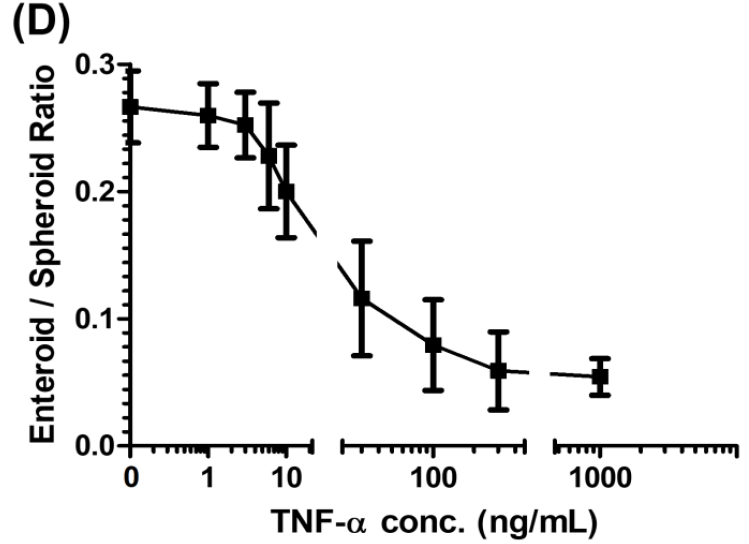
The response of intestinal organoid according to TNFα concentration in culture medium. (A) Morphological changes in control jejunal organoids with incremental increases in the TNFα concentration (n = 2). (B) Morphological changes in ileal organoids with incremental increases in the TNFα concentration (n = 2). (C) MTT assay for cell viability (n = 4). (D) Enteroid/spheroid ratio of human intestinal organoids (n = 4).

**Supplementary Figure S3.**
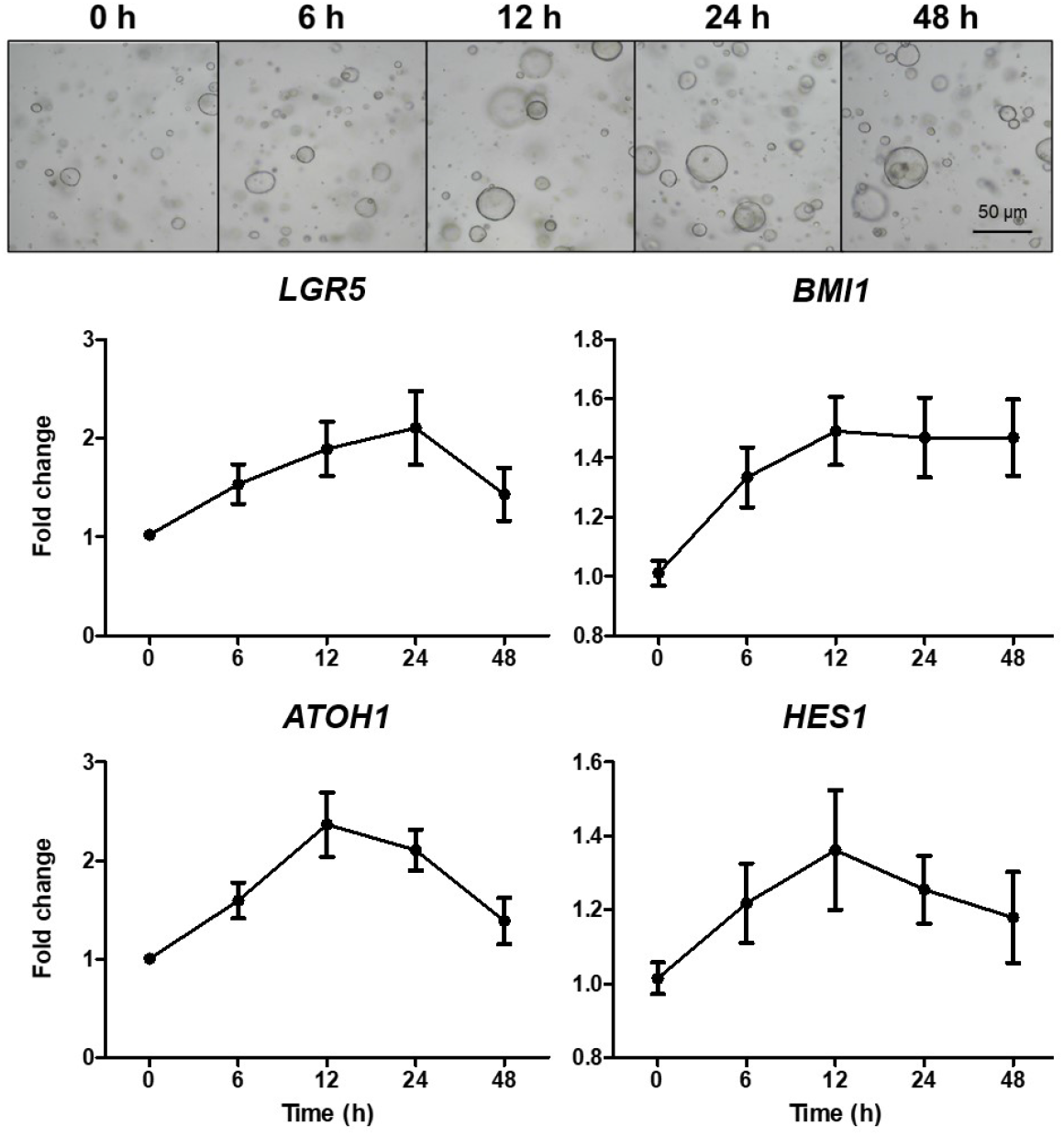
Changes in the expression of *LGR5, BMI1, ATHO1*, and *HES1* in control intestinal organoids 6, 12, 24, and 48 h after treatment with 30 ng/mL TNFα. Quantitative reverse transcription polymerase chain reaction was conducted in triplicate.

**Supplementary Figure S4.**
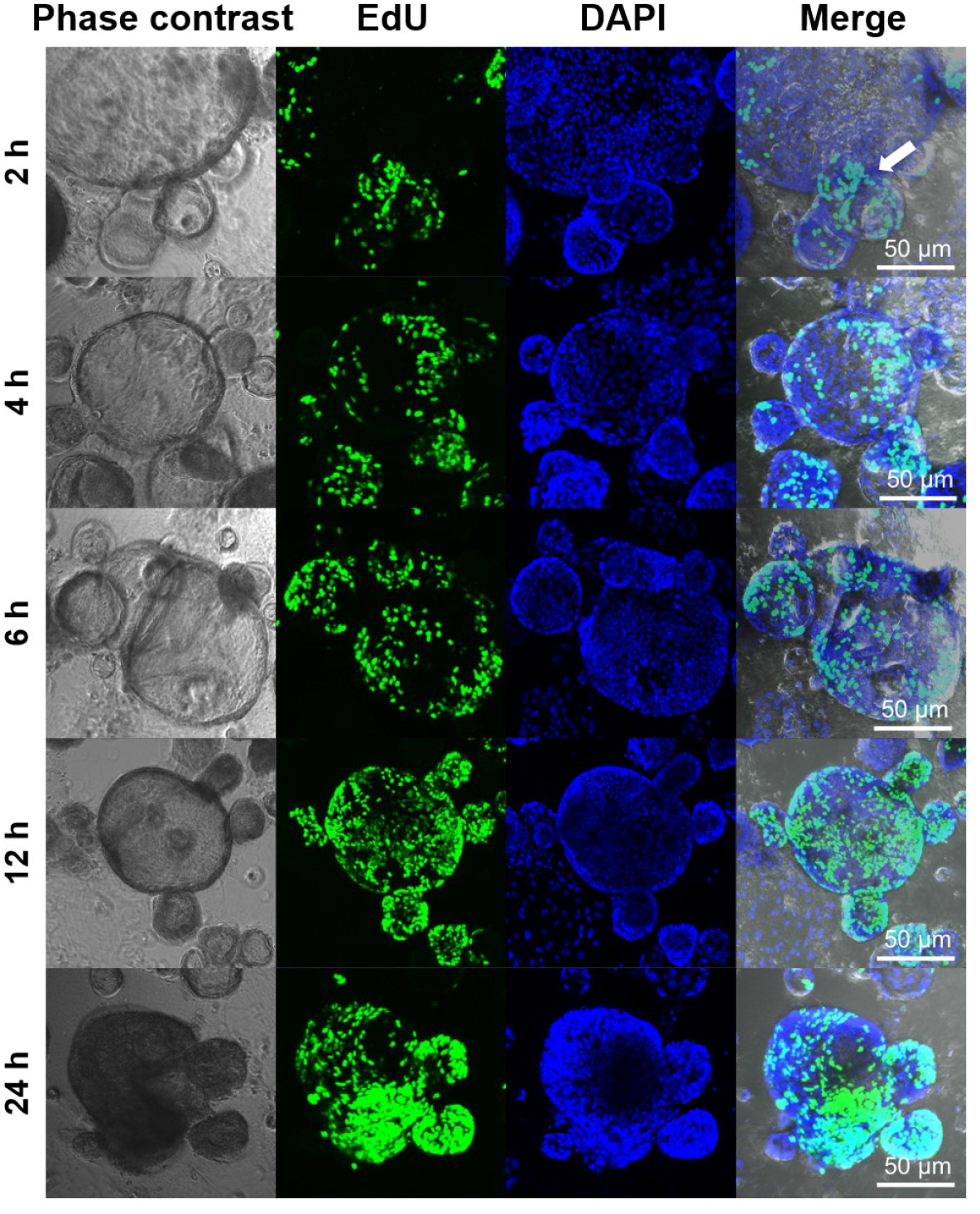
Distribution of EdU+ cells 2, 4, 6, 12, and 24 h after EdU administration in control organoid culture medium.

